# BPIFB3 facilitates flavivirus infection by controlling RETREG1-dependent reticulophagy

**DOI:** 10.1101/333435

**Authors:** Azia S. Evans, Nicholas J. Lennemann, Ka man Fan, Carolyn B. Coyne

## Abstract

The *flavivirus* genus, which includes dengue virus (DENV) and Zika virus (ZIKV), are significant human pathogens and the prevalence of infected vectors continues to geographically expand. Both DENV and ZIKV rely on expansion of the endoplasmic reticulum (ER) and the induction of autophagy to establish a productive viral infection. However, little is known regarding the interplay between the requirements for autophagy initiation during infection and the mechanisms used by these viruses to avoid clearance through the autophagic pathway. We recently showed that DENV and ZIKV inhibit reticulophagy (specific degradation of the ER through autophagy) by cleaving reticulophagy regulator 1 (RETREG1), an autophagy receptor responsible for targeted ER sheet degradation. These data suggest that DENV and ZIKV require specific autophagic pathways for their replication, while other autophagic pathways are antiviral. We previously identified BPI Fold Containing Family B Member 3 (BPIFB3) as a regulator of autophagy that negatively controls enterovirus replication. Here, we show that in contrast to enteroviruses, BPIFB3 functions as a positive regulator of DENV and ZIKV infection and that its RNAi-mediated silencing drastically inhibits the formation of viral replication organelles. We show that BPIFB3 depletion enhances ER fragmentation, while its overexpression protects against autophagy-induced ER degradation, demonstrating that BPIFB3 serves as a specific regulator of ER turnover. We further show that the antiviral effects of BPIFB3 depletion on flavivirus infection are reversed in RETREG1-depleted cells, and that BPIFB3 associates with RETREG1 within the ER, suggesting that BPIFB3 regulates a RETREG1-specific reticulophagy pathway. Collectively, these studies identify BPIFB3 as a regulator of the reticulophagy pathway and define the requirements for a novel host regulator of flavivirus infection.

**Author Summary:** Flaviviruses and other arthropod transmitted viruses represent a widespread global health problem with limited treatment options currently available. Thus, greater knowledge of the host factors required for replication and transmission is needed to provide a better understanding of the cellular requirements for infection. Here, we show that the endoplasmic reticulum (ER) localized protein, BPIFB3 is required to facilitate flavivirus infection. Depletion of BPIFB3 in cells inhibits dengue virus and Zika virus infection prior to replication of the viral genome. Mechanistically, we show that BPIFB3 inhibits ER degradation in an autophagy-specific manner and that loss of BPIFB3 decreases the availability of ER membranes needed for flavivirus replication. We further show that BPIFB3 specifically regulates the RETREG1 pathway, but not other pathways of ER turnover. Together, our data define a previously uncharacterized method of regulating ER degradation and show that BPIFB3 is an essential host factor for a productive flavivirus infection.

## Introduction

Flaviviruses, which include dengue virus (DENV) and Zika virus (ZIKV), are enveloped, positive-sense RNA viruses that replicate exclusively within endoplasmic reticulum (ER) membranes (1,2). Upon entry and uncoating, the flaviviral genome is directly translated as a single polyprotein and embedded in the ER, which induces ER expansion and the formation of viral replication organelles (3–6). Like other RNA viruses, flaviviruses sequester their replication machinery within membrane bound compartments that provide a high concentration of host and viral replication factors, while isolating viral replication intermediates from detection by the host innate immune system (7,8).

The success of flavivirus replication is closely linked to the availability of ER membranes. The ER is an expansive network of membranous sheets and tubules that originate at the nuclear envelope and extend to the cell periphery (9). ER sheets are distributed perinuclearly and function as the primary location of protein synthesis, while ER tubules extend to the plasma membrane and play a prominent role in lipid synthesis and communication with other organelles (10). Both DENV and ZIKV utilize rough ER as the primary source of membranes for the formation of replication organelles, while smooth ER is used in the formation of convoluted membranes during viral replication (7). These convoluted membranous formations are found in close proximity to replication complexes and mitochondria, suggesting a role in lipid synthesis and/or membrane expansion during infection (8).

Although autophagy commonly serves as a proviral pathway for RNA viruses, these viruses often avoid clearance by flux through the autophagic pathway, which can function in an antiviral manner (11). In the absence of infection, macroautophagy, hereafter referred to as autophagy, functions to degrade bulk cytoplasmic contents and excess or damaged organelles by fusion with the lysosome, which is referred to as autophagic flux (12,13). During infection, autophagy functions as an innate defense pathway to facilitate the clearance of viral complexes and infectious particles (14–16). Furthermore, recent evidence suggests that autophagy can be specifically activated through innate immune signaling to induce the clearance of intracellular pathogens (17). To overcome autophagy-mediated clearance, many viruses have developed strategies to either inhibit flux through the autophagic pathway or to utilize it in a proviral manner for virion maturation and release (14). However, the full relationship between flaviviruses and autophagy remains unclear. Evidence suggests that DENV and ZIKV induce autophagy as a mechanism to promote viral infection (18,19). Specifically, DENV infection promotes the degradation of lipid droplets by autophagy, termed lipophagy, to allow for an increased energy supply during infection (20,21). In contrast, we have previously demonstrated that degradation of the ER through selective autophagy (reticulophagy) restricts flavivirus replication due to the dependence of viral replication on ER membranes (22). Furthermore, DENV and ZIKV specifically inhibit reticulophagy by cleaving reticulophagy regulator 1 (RETREG1), which is necessary for targeted ER sheet degradation (22). These studies collectively suggest that specific autophagic pathways may be anti-flaviviral, while others are pro-flaviviral.

We previously identified bactericidal/permeability increasing protein (BPI) fold containing family B member 3 (BPIFB3) as an ER-localized antiviral regulator of coxsackievirus B (CVB)infection, a positive-sense RNA virus belonging to the *Enterovirus* genus, through its negative regulation of a non-canonical form of autophagy (23). Similar to flaviviruses, CVB relies on the availability of intracellular membranes to establish replication compartments; however, the source of these cellular membranes is variable. In this study, we identified BPIFB3 as a positive regulator for flavivirus infection. Mechanistically, we show that BPIFB3 functions upstream of RETREG1 to specifically control reticulophagy, thereby controlling the availability of ER membranes for viral replication. Our study therefore defines a specific role for BPIFB3 in reticulophagy and suggests that it differentially controls enterovirus and flavivirus replication.

## Results

### BPIFB3 is required for DENV and ZIKV infection

We have previously shown that BPIFB3 is an ER-localized negative regulator of CVB infection (23). Given that flaviviruses replicate exclusively on membranes derived from the ER, we determined whether BPIFB3 also functions to regulate DENV and ZIKV replication. To do this, human brain microvascular endothelial cells (HBMEC) were transfected with either a siRNA targeting BPIFB3 (BPIFB3si) or a control siRNA (CONsi) and infected with DENV, ZIKV, or CVB (23). In contrast to the significant enhancement of CVB infection, RNAi-mediated silencing of BPIFB3 resulted in an approximately 90% decrease in the replication of both DENV and ZIKV, as assessed by RT-qPCR and immunofluorescence microscopy for the production of double-stranded RNA, a replication intermediate (**Figure 1a, d, S1d**), and a 100-fold decrease in infectious particle production (**Figure 1b**). This phenotype was specific for BPIFB3 and other members of the BPIFB family (BPIFB2 and BPIFB6) did not affect DENV and ZIKV uniformly (**Figure S1a, b)**. Depletion of BPIFB3 was confirmed by RT-qPCR (**Figure S1c**). To determine at which stage of the flaviviral life cycle BPIFB3 depletion impairs, we used HBMEC stably propagating a DENV subgenomic replicon (HBMEC^rep^) (24). DENV replicon cells express the full seven nonstructural proteins that allows for replication of replicon RNA as well as the remodeling of ER membranes, including the formation of replication organelles, similar to viral infection (25). Silencing of BPIFB3 in HBMEC^rep^ had no effect on replicon RNA levels (**Figure 1c**), suggesting the defect in flavivirus infection occurs prior to the formation of viral replication organelles within the ER.

**Figure 1.**
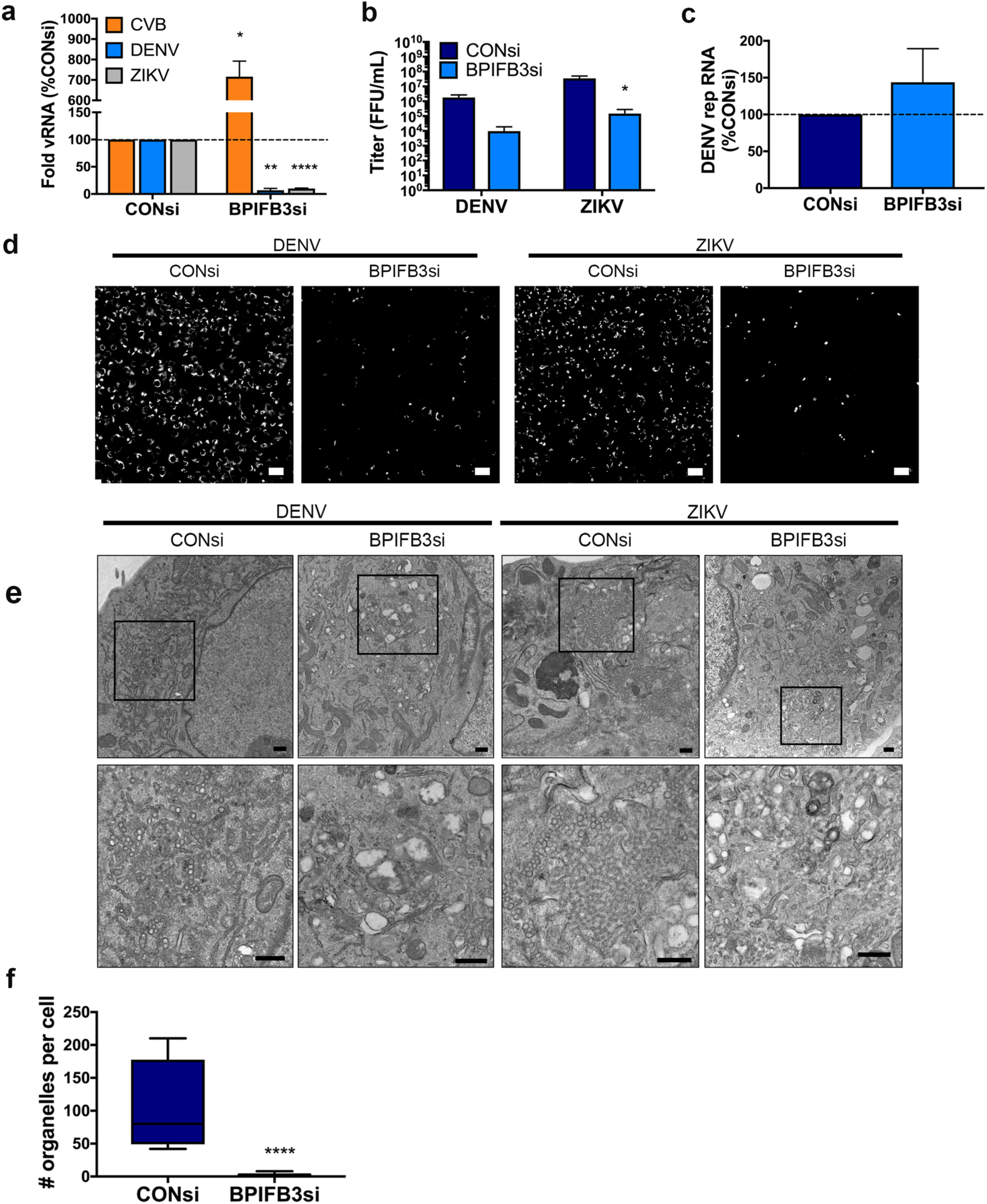
BPIFB3 depletion restricts an early step in flavivirus infection. **(a)** Infection levels of CVB, DENV, and ZIKV determined by RT-qPCR. Data are presented as a percent change from CONsi-transfected cells. **(b)** Titration (by fluorescence focus unit assay) of DENV and ZIKV infectious particle production from HBMEC from panel (a). **(c)** DENV replicon RNA levels as determined by RT-qPCR in response to BPIFB3 depletion, presented as percent of CONsi. **(d)** Immunofluorescence microscopy for dsRNA (green), a replication intermediate, in CONsi or BPIFB3si transfected HBMEC. Scale bar is 50 μm. **(e)** TEM from HBMEC transfected with CONsi or BPIFB3si and infected with DENV or ZIKV. Top panel scale bar is 2 μm and bottom panel scale bar is 500 nm. **(f)** Quantification of the number of ZIKV replication organelles per cell in TEM images (panel e). Students t test were performed to determine statistical significance (*< 0.05, **< 0.01, *** < 0.001, **** < 0.0001).

To characterize the effects of BPIFB3 depletion on DENV and ZIKV infection, we performed transmission electron microscopy (TEM) on DENV- and ZIKV-infected HBMEC transfected with CONsi or BPIFB3si. We found that BPIFB3si prevented the formation of both DENV and ZIKV membrane bound viral replication organelles (**Figure 1e**). Quantification of ZIKV replication organelles (defined as ER associated, membrane bound vesicles of approximately 70-100 nm) confirmed that BPIFB3 silencing inhibited flavivirus infection prior to genome replication (**Figure 1f**). In addition to defects in replication organelle formation, we did not observe the formation of convoluted membranes or ER rearrangement characteristic of flavivirus infection in BPIFB3-depleted cells, suggesting BPIFB3si inhibits infection early during the viral lifecycle.

### Infection of BPIFB3 depleted cells induces aberrant ER structures

We showed previously that BPIFB3 localizes to domains enriched for the ER sheet marker CLIMP63 (26). Therefore, we sought to examine whether BPIFB3 is involved in regulating ER morphology or turnover during flavivirus infection. In uninfected cells, ER sheets originate at the nuclear envelope and extend to the cell periphery in a fairly uniform arrangement; however, during infection with DENV and ZIKV, ER sheets (marked by CLIMP63) condense around the perimeter of the nucleus where they co-localize with viral double stranded RNA (dsRNA), designating the location of viral membrane remodeling and replication organelle formation (22). In some cases, cells depleted of BPIFB3 exhibited low levels of viral replication (as determined by dsRNA immunofluorescence); however, these cells exhibited an abnormal rearrangement of CLIMP63-positive ER domains into punctate structures (**Figure 2a**). The ability to establish infection in select BPIFB3si cells may be caused by variations in knockdown efficiency across individual cells,or could indicate that in some cases, low levels of replication can be initiated in cells depleted of BPIFB3. TEM analysis of BPIFB3 depleted cells infected with ZIKV exhibited dramatic expansions of ER membranes with no evidence of replication organelles (**Figure 2b**). Importantly, uninfected BPIFB3si transfected cells did not exhibit aberrant ER structures and contained few identifiable ER membranes; suggesting that ZIKV may be able to infect BPIFB3 depleted cells and induce early ER remodeling but is unable to readily form replication organelles or establish efficient replication.

**Figure 2.**
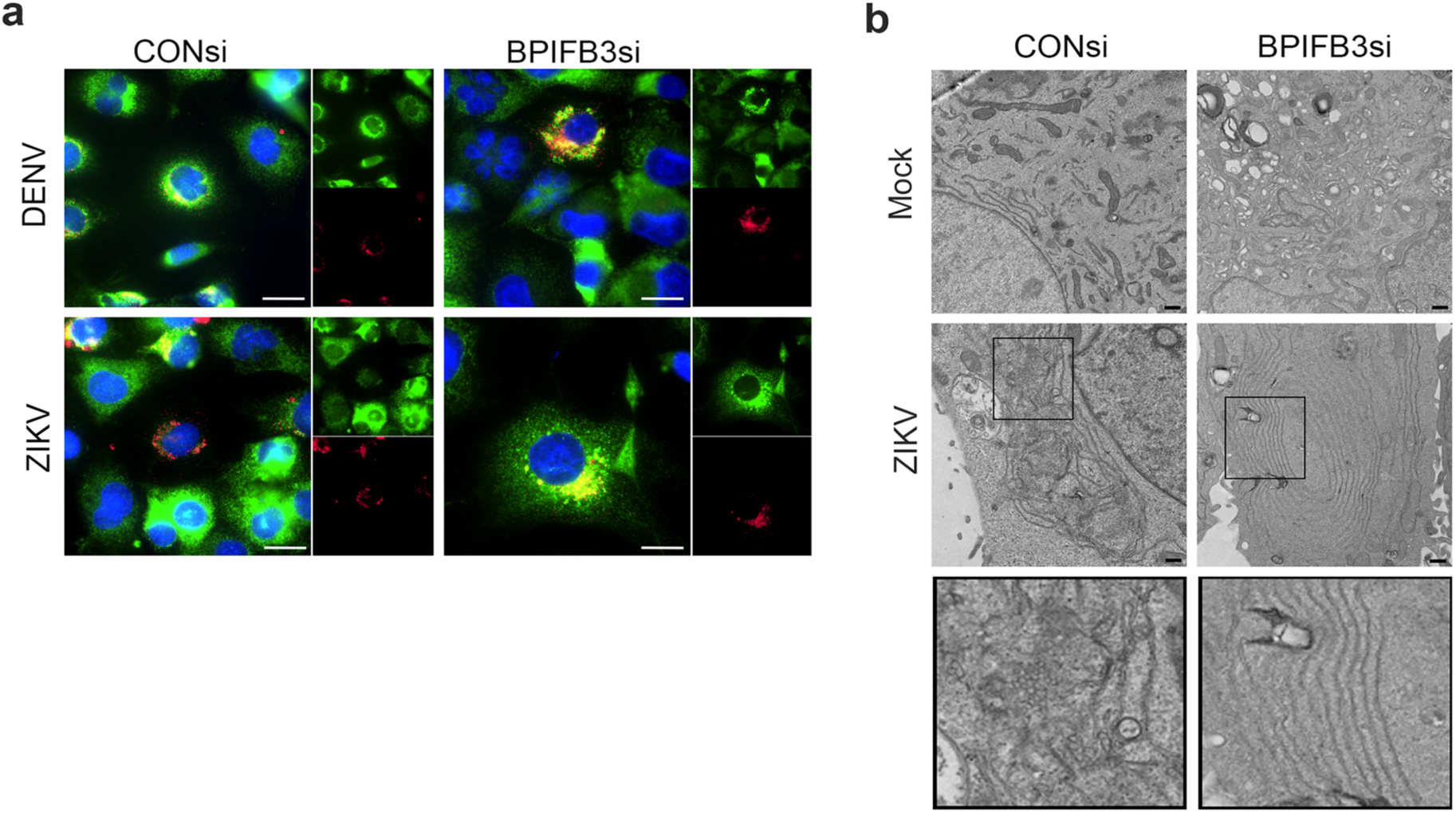
BPIFB3 depletion induces aberrant ER structures during flavivirus infection. **(a)** Immunofluoresence microscopy from CONsi or BPIFB3si transfected HBMEC infected with DENV or ZIKV (MOI=1) and stained for dsRNA (red) and Climp63 (green) 48hrs post-infection. Scale bars are 20 μm. **(b)** TEM images from CONsi or BPIFB3si transfected HBMEC infection with ZIKV (or mock infected controls). Scale bars are 2 μm.

### BPIFB3 regulates ER sheet morphology in response to autophagy induction

We previously demonstrated that BPIFB3 serves as a regulator of a non-canonical form of autophagy, however the precise mechanism of regulation remained unclear (23). To assess the impact of BPIFB3 silencing on host cell pathways, we performed whole transcriptome RNAseq studies on uninfected and infected HBMEC transfected with CONsi or BPIFB3si. Differential expression analysis identified numerous autophagy related genes as dysregulated in BPIFB3si samples compared to controls (**Figure 3a**). These included ATG101, WIPI3, ATG3, ATG12, and LC3B which were transcriptionally upregulated in BPIFB3 depleted cells and are involved in autophagosome formation prior to vesicle release and maturation (12). Interestingly, WIPI2, BECN1, ATG10, and ATG7 were downregulated in BPIFB3si cells despite their function during autophagosome formation in conjunction with the upregulated genes mentioned above. These results further suggested that BPIFB3 is involved in regulating a non-canonical form of autophagy. Importantly, our transcriptional analyses did not reveal an increase in interferons (IFNs) or interferon stimulated gene (ISG) production in uninfected or infected samples, confirming that the decrease in infection observed in BPIFB3 depleted cells is not the result of increased innate immune activation or signaling. We also examined the expression levels of ER structural transcripts and found that the ER sheet marker CLIMP63 (also named CKAP4) was upregulated in uninfected BPIFB3si samples, while the ER tubule protein reticulon3 (RTN3) was downregulated. These transcriptional changes may suggest a need for increased ER sheet production in response to BPIFB3 silencing.

**Figure 3.**
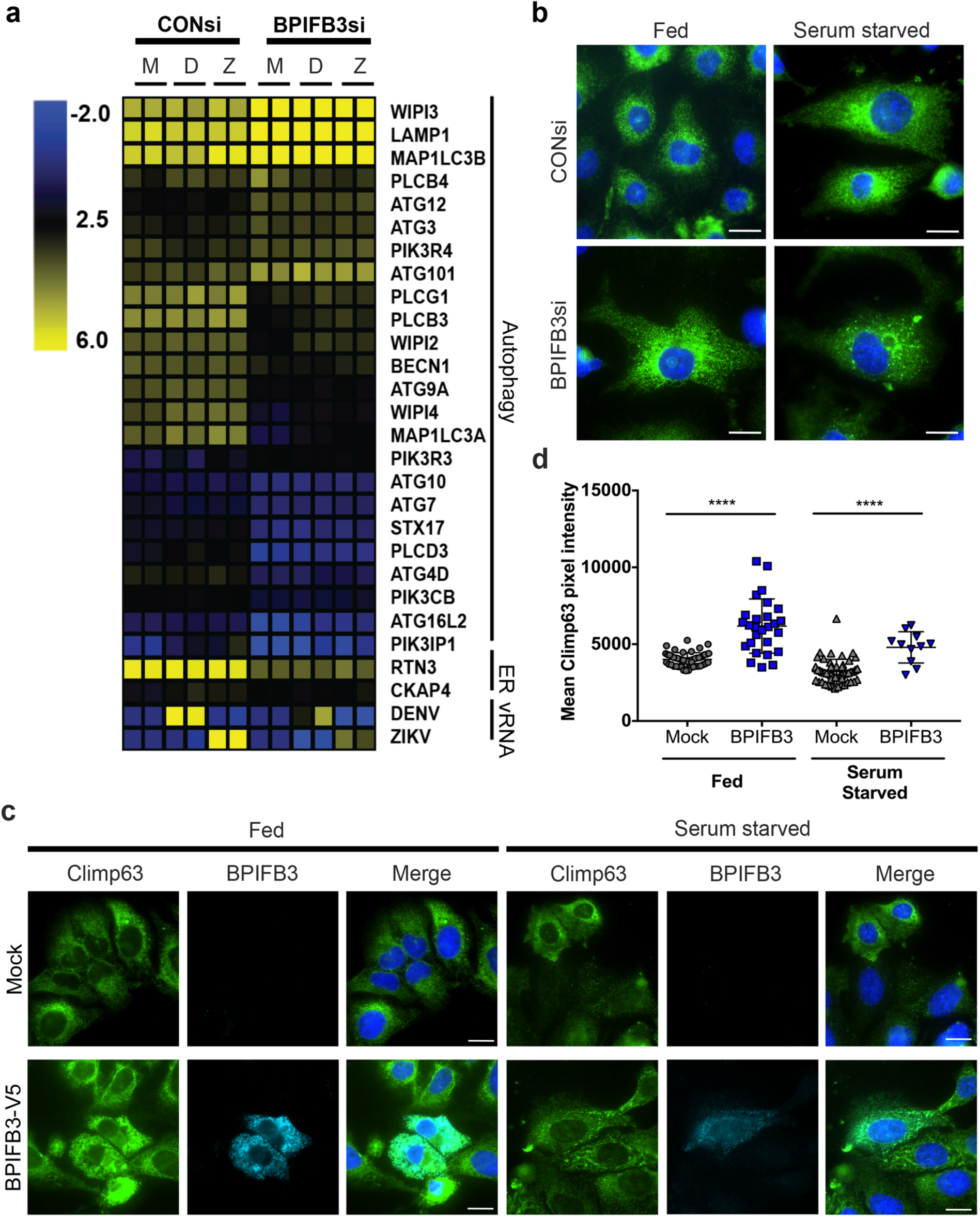
BPIFB3 controls ER turnover through an autophagy-associated pathway. **(a)** Heatmap of log_2_(RPKM) values as determined by RNASEq from mock (M), DENV (D), or ZIKV (Z)-infected CONsi and BPIFB3si transfected HBMEC. **(b)** Climp63 (green) ER morphology of CONsi and BPIFB3si HBMECs under fed and serum starved conditions. **(c)** U2OS cells expressing BPIFB3-V5 or Mock transfected were either fed or serum starved and stained for Climp63 (green) and V5 (teal). **(d)** Quantification of Climp63 pixel intensity from panel c shows BPIFB3 expression corresponds to increased levels of the ER sheet marker. All scale bars are 20 μm. A non-parametric Kruskal-Wallis test was performed to determine significance of IF pixel quantification (**** < 0.0001).

Given that we observed an enhancement in ER-enriched punctae in DENV and ZIKV infected BPIFB3si cells (**Figure 2a**), we next determined whether BPIFB3 enhanced ER degradation through autophagy. To do this, we examined ER sheet morphology by immunofluoresence under both nutrient rich (fed) or serum starved conditions, which induces autophagy. We found that serum starvation of cells depleted of BPIFB3 induced the formation of CLIMP63 puncta that were absent in CONsi transfected cells (**Figure 3b**). To determine if this was unique to ER sheets, or if BPIFB3si also influenced the rearrangement of ER tubules, we analyzed the localization of the tubule specific protein reticulon4 (RTN4). Serum starvation induced the rearrangement of RTN4 positive ER tubules, however this phenotype was also observed in CONsi cells, and was not further exaggerated by BPIFB3 depletion (**Figure S2**).

To determine if ectopic expression of BPIFB3 exerted an opposing phenotype leading to the stabilization of ER sheets, we transfected human osteosarcoma U2OS cells with V5 fused BPIFB3 (BPIFB3-V5) and transferred them to either nutrient rich media (fed) or HBSS (serum starved) 48 hours post transfection. Using quantitative image analysis, we found that the levels of endogenous CLIMP63 were significantly higher in cells overexpressing BPIFB3 under both conditions (**Figure 3c, 3d**). These data suggest that BPIFB3 expression protects ER sheets from degradation through autophagy, and that loss of BPIFB3 leads to enhanced ER turnover.

### RETREG1 and BPIFB3 localize in close proximity within the ER

Given our data indicating BPIFB3 stabilizes ER sheets, we sought to determine whether BPIFB3 regulates the reticulophagy pathway. We first analyzed whether BPFIB3 localizes with RETREG1, which we showed previously functions as an antiviral regulator of flavivirus infection (22).We found that ectopically expressed BPIFB3 colocalized with both wild-type RETREG-1 and an autophagy deficient mutant of RETREG1 lacking an LC3 interacting region (LIR) (mutLIR), suggesting that BPIFB3 and RETREG1 colocalize independent of the ability of RETREG-1 to function in reticulophagy (**Figure 4a**). To assess if co-localization was due to a direct interaction we performed co-immunoprecipitation experiments, however we were unable to observe an interaction between BPIFB3 and RETREG1 due to low protein solubility.

**Figure 4.**
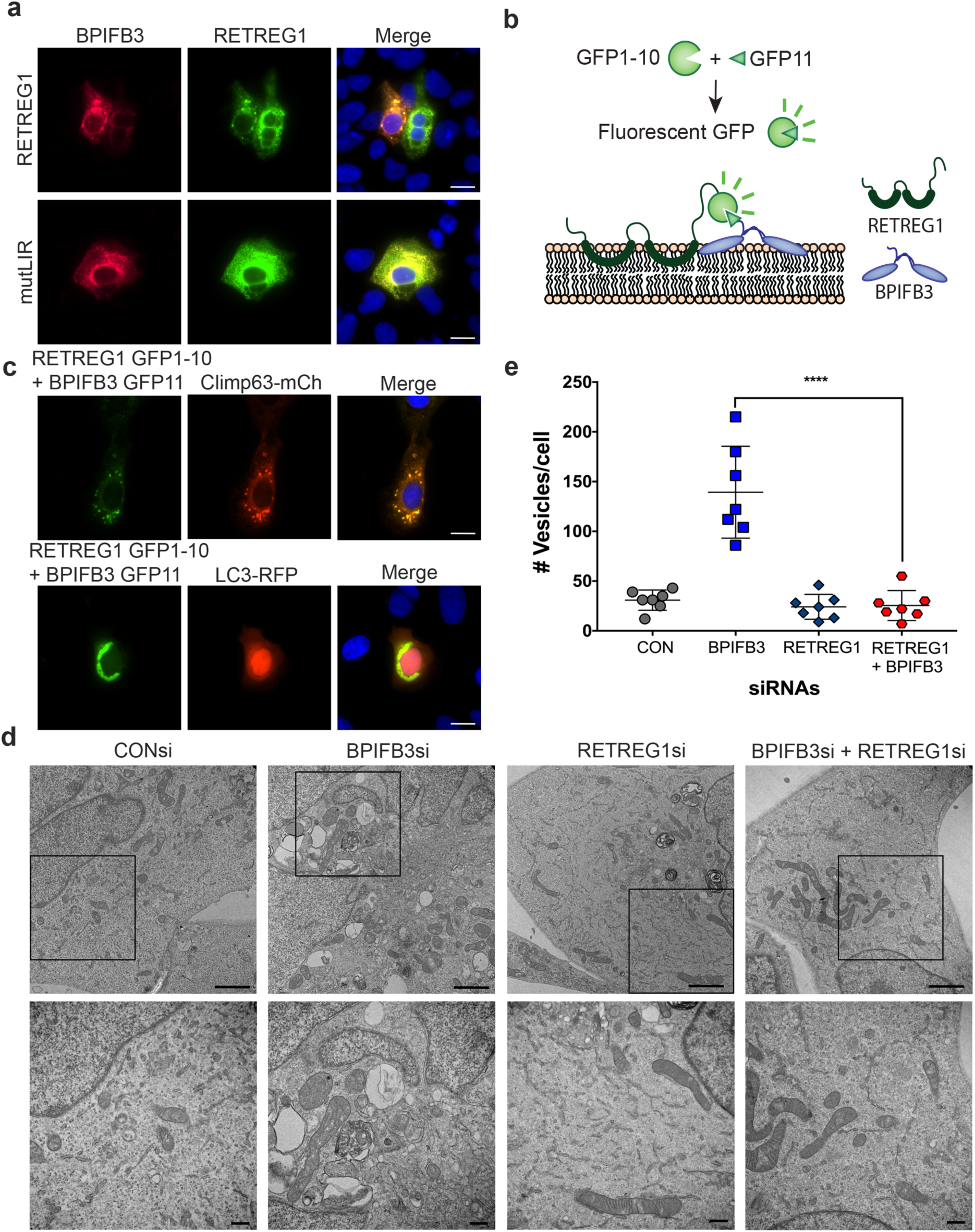
BPIFB3 and RETREG1 co-ocalize to ER sheets. **(a)** U2OS cells transfected with BPIFB3-V5 and RETREG1-GFP or RETREG1 mutLIR GFP. **(b)** Schematic of modified BiFC assay using split GFP. **(c)** Split GFP assay, RETREG1 GFP1-10 and BPIFB3 GFP11 were co-expressed with Climp63-mCherry or LC3-RFP. Green fluorescence indicates RETREG1 and BPIFB3 are in close enough proximity to interact. Immunofluorescence scale bars are 20 μm. **(d)** TEM of HBMEC depleted of BPIFB3 and RETREG1 alone or together. Top panel shows total cell morphology, scale bars represent 2 μm. Black boxes indicated regions magnified in bottom panel to show ER membrane and vesicle morphology. Lower panel scale bars are 500 nm. **(e)** Quantification of total cytoplasmic vesicles from seven cells in each knockdown condition. A One-way ANOVA was performed to determine significance (**** < 0.0001).

To determine whether BPIFB3 and RETREG1 reside in close proximity to one another, we used a modified reversible bimolecular fluorescence complementation (BiFC) assay. This assay utilizes GFP broken into two distinct fragments, a large portion composed of the first ten β sheets of GFP (GFP1-10) and a smaller fragment composed of the eleventh β sheet (GFP11) (27). When the tagged proteins do not interact or associate within the same complex, the GFP fragments are too far apart and there is no fluorescence. However, if there is either a direct or indirect association (of less than 10nm apart), GFP folds correctly and fluoresces similar to full length GFP (28) (schematic, **Figure 4b**). We fused RETREG1 with GFP1-10 and BPIFB3 with the smaller GFP11. U2OS cells were transfected with each split GFP construct and with either mCherry fused CLIMP63 or RFP fused LC3. We found that RETREG1 GFP1-10 and BPIFB3 GFP11 localized in very close proximity (<10nm) to one another, as determined by positive GFP fluorescence in cotransfected cells, however no green fluorescence was observed when RETREG1 GFP1-10 was expressed with GFP11 alone (**Figure S3**). We further found that BiFC-component BPIFB3 and RETREG1 co-localized with the ER sheet marker CLIMP63, but not the autophagosome marker LC3 (**Figure 4c**). These data suggest that BPIFB3 and RETREG1 localize to ER sheets and are in close proximity, but may not form a direct interaction. Further, we did not observe RETREG1 colocalization with LC3B upon expression of BPIFB3, thus BPIFB3 may restrict RETREG1-mediated reticulophagy.

### BPIFB3 negatively regulates RETREG1-mediated reticulophagy

To determine whether silencing of BPIFB3 enhances reticulophagy, we first analyzed ER morphology and autophagosome accumulation by TEM in cells co-depleted of BPIFB3 and RETREG1. We previously showed that BPIFB3 silencing leads to an accumulation of autophagosomes, lysosomes, and amphisomes (23). However, this phenotype was completely reversed in cells transfected with RETREG1si (**Figure 4d, 4e**), suggesting that this induction occurs downstream of RETREG-1-mediated reticulophagy.

In response to autophagy induction, RETREG1 interacts with LC3 to target ER membranes to the autophagosome for eventual degradation by the lysosome (13). Given that BPIFB3 expression led to increased levels of ER sheets and that silencing of RETREG1 reversed the induction of autophagy in cells transfected with BPIFB3si, our data suggested that BPIFB3 inhibits RETREG1-mediated reticulophagy. To confirm this, we transfected U2OS cells with RETREG1-GFP and LC3-RFP in the presence or absence of BPIFB3 under nutrient rich or nutrient deprived conditions. We found no differences in the numbers of RETREG1 positive puncta alone (**Figure 5b**) or in the co-localization of RETREG1 and LC3 puncta (**Figure 5c**) under nutrient rich conditions. However, in response to nutrient deprivation, we found that BPIFB3 expression significantly reduced the number or RETREG1 positive puncta (**Figure 5b**) and prevented the co-localization of RETREG1 with LC3 (**Figure 5c**). Collectively, these data suggest that BPIFB3 functions as a negative regulator of RETREG1-mediated reticulophagy.

**Figure 5.**
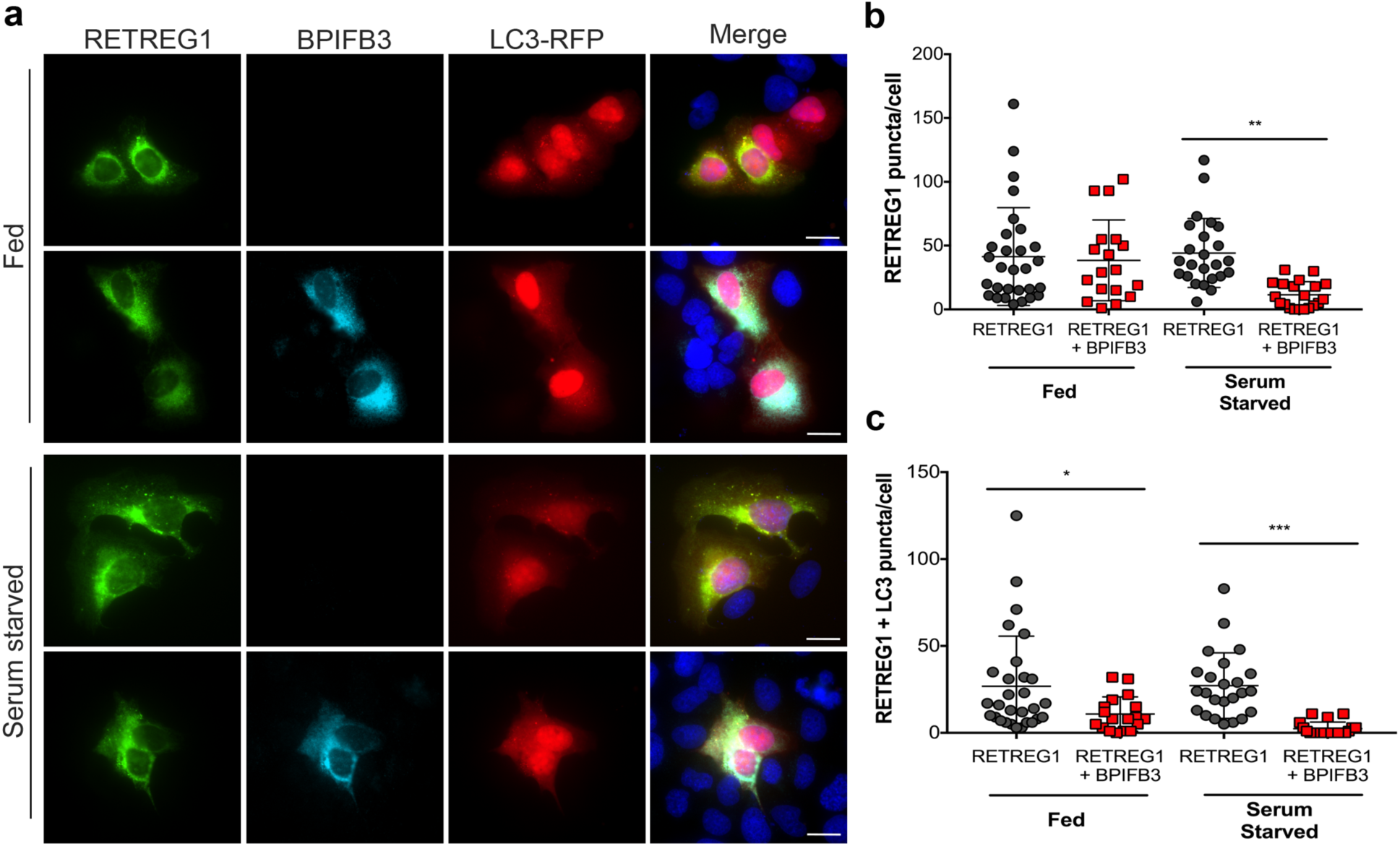
BPIFB3 inhibits RETREG1 reticulophagy in response to nutrient deprivation. **(a)** U2OS cells co-expressing RETREG1-GFP and LC3-RFP were co-transfected with or without BPIFB3-V5 and kept under fed or serum starved conditions. **(b)** Total RETREG1 puncta were quantified under each condition as indicated. **(c)** Quantification of the number of RETREG1 and LC3 positive positive puncta under either nutrient rich (fed) or nutrient deprived (serum starved) conditions. A One-way ANOVA with Bonferroni correction was performed to determine significance (* < 0.05, ** < 0.01, *** < 0.001).

### BPIFB3 facilitates flavivirus replication by negatively regulating reticulophagy

DENV and ZIKV are dependent on the availability of ER membranes to replicate and reticulophagy thus functions as an antiviral pathway that limits the availability of these membranes (22). To determine if the reduction of flavivirus replication in cells depleted of BPIFB3 resulted from enhanced reticulophagy, we co-depleted BPIFB3 and RETREG1 in cell and infected with DENV or ZIKV. We found that silencing of RETREG1 completely reversed the inhibition of flavivirus infection in cells silenced for BPIFB3 expression, as determined by both qPCR for vRNA and FFU for viral titers (**Figure 6a, 6b**). Consistent with our previous work (22), silencing of RETREG1 enhanced flavivirus replication, which was unaffected by BPIFB3 silencing. Moreover, we found that co-depletion of BPIFB3 and RETREG1 also reversed the proviral impact of BPIFB3 silencing on CVB replication (**Figure 6c**). Depletion of BPIFB3 and RETREG1 was confirmed by RT-qPCR (**Figure S4**). Interestingly, we found that BPIFB3 specifically regulates RETREG1-mediated ER sheet reticulopahgy as the anti-flaviviral effect of BPIFB3 silencing was unaffected by silencing of either reticulon 3 (RTN3), the ER tubule specific reticulophagy receptor, or the ER stress-specific reticulophagy receptor Sec62 (**Figure S5**). Lastly, we found that BPIFB3 localizes to the same ER domains as flaviviral nonstructural proteins during active replication, as determined by the colocalization of DENV NS1, NS3 and ZIKV NS4B which localize to replication organelles during infection (**Figure 6d**). These data implicate BPIFB3 as a specific negative regulator of RETREG1-mediated reticulophagy, which functions to promote flaviviral replication.

**Figure 6.**
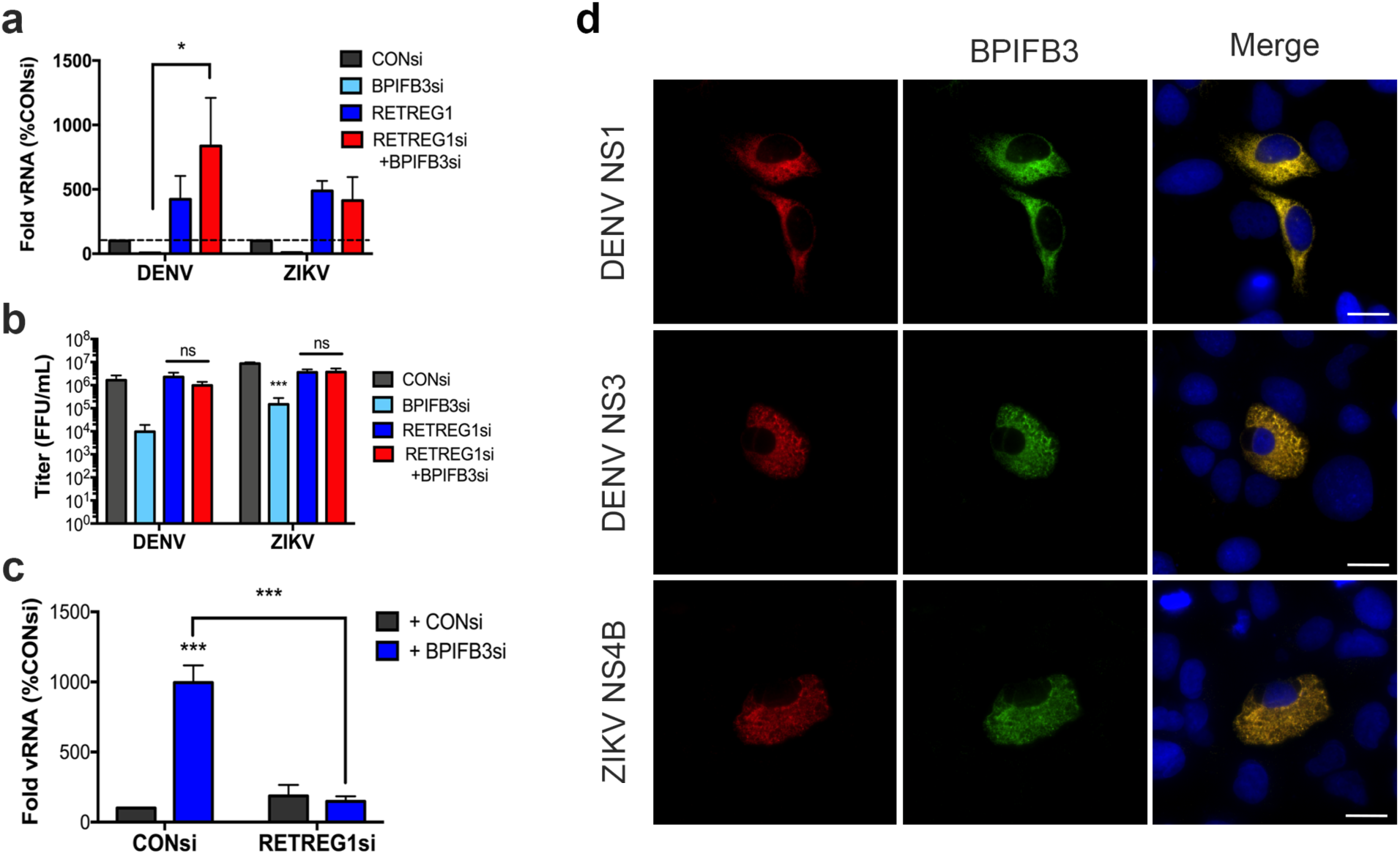
BPIFB3 regulates RETREG1 specific reticulophagy during flavivirus infection. **(a)** RT-qPCR for DENV and ZIKV infection levels in cells depleted of BPIFB3 and RETREG1 alone or together. **(b)** Infectious particle production from BPIFB3 and RETREG1 depleted cells as determined by fluorescence focus unit (FFU) assays. **(c)** Cells depleted of BPIFB3 and RETREG1 were infected with CVB at a MOI of 1 and analyzed for level of infection by RT-qPCR. Statistical significance was determined by 2-way ANOVA for each panel (* < 0.05, *** < 0.001). **(d)** U2OS cells overexpressing DC-SIGN to enhance infection were transfected with BPIFB3-V5 (green) and then infected with DENV or ZIKV. Cells were fixed and stained 48hrs following infection and immunostained for the indicated viral proteins (in red).

### Discussion

The success of flavivirus infection depends on the cooperation of numerous cellular organelles and pathways that function to produce progeny virions, specifically relying on host membranes throughout their lifecycles. Here we show that BPIFB3 is required for DENV and ZIKV infection by regulating the availability of ER membranes for viral remodeling. Our data show that BPIFB3 depletion enhances ER sheet reticulophagy in response to viral infection and the induction of autophagy. Furthermore, we demonstrate BPIFB3 functions to regulate RETREG1 targeted reticulophagy and not RTN3 or Sec62 specific pathways. These findings thus not only define the role of specific autophagic pathways in the regulation of flavivirus infection, but also identify BPIFB3 as a novel regulator of RETREG1-specific forms of reticulophagy.

Unlike other RNA viruses, flaviviruses depend solely on ER-derived membranes for their replication. The viral genome is delivered to the rough ER following entry and uncoating, where translation of viral proteins induces expansion of the ER. Of the seven nonstructural proteins, the majority remain associated with the ER throughout the lifecycle, where they function in viral replication, membrane remodeling, and inactivation of reticulophagy and ER stress pathways (5–7,22,29,30). While it has been suggested that the virally-encoded non-structural proteins NS1, NS4A, and NS4B are involved in membrane manipulation during DENV infection, little is known regarding host factors essential for this process. Currently, only three host factors have been implicated in membrane expansion, including fatty acid synthase, RETREG1, and reticulon 3.1A (RTN3.1A). FASN is recruited to sites of replication organelle formation by the DENV protease NS3, suggesting increased lipid synthesis is important for membrane remodeling (20). Additionally, both DENV and ZIKV inhibit ER degradation by cleaving the RETREG1 reticulophagy receptor, allowing for an accumulation of ER membranes (22). Lastly, RTN3.1A localizes to viral replication organelles to facilitate proper membrane curvature, however it does not interact with DENV or ZIKV NS4a during membrane remodeling (31). Our work presented here further confirms that degradation of the ER is an antiviral process and defines a new mechanism used by flaviviruses to regulate ER turnover. RNAi mediated silencing of BPIFB3 leads to enhanced levels of reticulophagy, which decreases the availability of ER membranes for flavivirus replication. Concurrent depletion of RETREG1 with BPIFB3 overcomes this defect, demonstrating that the antiviral effects of BPIFB3 depletion are specific to RETREG1-mediated reticulophagy and inhibition of this pathway restores viral replication. These data imply that the manipulation of BPIFB3 protein levels during flavivirus infection could alter reticulophagy levels to either enhance viral replication or allow for the host cell to overcome infection at an early stage.

One method proposed to promote membrane expansion during flavivirus infection is the induction of autophagy (30). However, our data demonstrate that enhanced levels of reticulophagy, particularly early during infection, inhibits membrane remodeling and replication organelle formation. Recent work has identified a number of ER-specific autophagy pathways that differ by the receptor used to target cargo to autophagosomes (13,32,33). However, it remains unclear whether these pathways are regulated by the same machinery that controls canonical macroautophagy. The growing diversity in the various forms of autophagy further complicates our understanding of the relationship between viral infection and this pathway, as certain forms of autophagy may differentially regulate viral replication at various stages of the viral life cycle. The work presented here, in combination with our previous work characterizing BPIFB3 as a negative regulator of CVB infection, demonstrates the unique requirements for autophagy between different RNA virus families. In contrast to the unclear role for distinct autophagic pathways in flavivirus infection, CVB benefits from autophagy induction, as it uses autophagosomes and other cytoplasmic vesicles for replication organelle formation. Importantly, CVB inhibits fusion of the autophagosome with the lysosome, which enhances the number of cytoplasmic vesicles and prevents the degradation of viral replication machinery (34–36). Conversely, it has not been demonstrated whether flaviviruses have developed strategies to avoid clearance through the macroautophagy pathway similar to CVB and other enteroviruses. While the induction of autophagy during flavivirus infection has been implicated in enhancing viral replication (37), the precise timing of induction may have distinct effects on the viral lifecycle. Furthermore, the ability to specifically activate one form of autophagy while inhibiting others may be essential for successful flavivirus infection. The distinction between membrane manipulation during CVB infection and flavivirus infection explains the differential effects of BPIFB3 in regulating these unique viruses and further suggests that increased flux through autophagy is detrimental to flavivirus replication.

The BPIFB family of proteins were initially named and identified because of their homology to the bactericidal/permeability-increasing (BPI) protein; a secreted antimicrobial protein that functions through binding to LPS (38–40). Despite the high degree of predicted structural homology, BPIFB3 localizes to the ER and is not secreted (23). Of the other members of the family, BPIFB2 and BPIFB6 are also ER localized, however neither appear to regulate autophagy (26) or flavivirus infection. BPIFB proteins contain two BPI folds demonstrated to have lipid binding properties. Unlike other BPIFB proteins, the first BPI domain (BPI1) of BPIFB3 lacks the ability to bind lipids, while BPI2 is capable of binding phosphatidic acid, phosphatidyserine, cardiolipin, and other lipid molecules (26). Of the related proteins, BPIFB6 is the only protein to be characterized, and has been demonstrated to regulate secretory trafficking and Golgi morphology (26). Together with the data presented here, this suggests that a possible unifying function of these proteins is to regulate sites of vesicle trafficking. Here we show BPIFB3 over expression decreases the amount of ER specific autophagosomes in a cell, while depletion enhances reticulophagy. In comparison, BPIFB6 depletion results in Golgi dispersal and a disruption of retrograde and anterograde trafficking (26). This alludes to a possible mechanism where BPIFB3 and BPIFB6 expression is associated with decreased vesicle trafficking to the autophagic and secretory pathways respectively, while loss of expression leads to enhanced vesicle trafficking originating in the ER. Importantly, expression of BPIFB3 is remarkably low, and we are unable to detect endogenous protein by either western or immunofluorescence. Despite its low expression, depletion of BPIFB3 elicits a dramatic phenotype in cells, suggesting an essential role in regulating morphology of the cellular membrane network. This is consistent with other ER structural proteins that drastically effect membrane morphology at very low levels of endogenous expression (41). Their potential roles in vesicle trafficking has important implications for the ability of these proteins to impact the trafficking and spread of a variety of viruses. However, further characterization is required to delineate the different methods by which viruses are trafficked during infection.

The relationship between flavivirus infection and the autophagic pathway is likely to be complex. While the initiation of autophagy and lipophagy have been demonstrated as proviral pathways (19,21,30,37), flux through the autophagic pathway and reticulophagy are antiviral (18,22,42). Thus, further characterization of the role of specific autophagic pathways in the regulation of flavivirus infection is needed to understand and develop new mechanisms to control infection.

## Methods

### Cells and viruses

Human brain microvascular endothelial cells (HBMEC) were maintained in in RPMI 1640 supplemented with 10% fetal bovine serum (FBS), 10% NuSerum, 1x non-essential amino acids, 1x minimum essential medium vitamins, 1% sodium pyruvate, and 1% antibiotic. Human bone osteosarcoma U2OS and Vero cells were grown in DMEM with 10% FBS and 1% antibiotic. Development of DENV replicon HBMECs using constructs provided by Theodore Pierson (NIH/NIAID) was described previously(24). *Aedes albopictus* midgut C6/36 cells were cultured in DMEM supplemented with 10% FBS and 1% antibiotic at 28°C in a 5% CO_2_ atmosphere.

DENV2 16881 and ZIKV Paraiba/2015 (provided by David Watkins, University of Miami)were propagated in C6/36 or Vero cells, respectively(43). Titers were determined by fluorescent focus assay as previously described, using recombinant anti-double-stranded RNA monoclonal antibody (provided by Abraham Brass, University of Massachusetts)(44). Propagation and titration have been describe previously of CVB3 (RD) has been described previously(45). Experiments measuring infection levels were performed using a multiplicity of infection (MOI) of 1 for 16 hours (CVB) or 48 hours (DENV and ZIKV), and infection was quantified by RT-qPCR or fluorescent focus assay.

### siRNAs, plasmids and transfections

Characterization of siRNAs targeting BPIFB3, BPIFB2, BPIFB6, and RETREG1 (FAM134B) have been described previously(22,23). Sequences of siRNAs targeting RTN3 or Sec62 are, RTN3: CCACUCAGUCCCAUUCCAUtt, and Sec62: GAAGGAUGAGAAAUCUGAAtt. All siRNAs, including the scrambled control (CONsi), were purchased from Sigma. Efficiency of knockdown was determined by RT-qPCR for each siRNA target. siRNAs were reverse transfected at 25 nM in to HBMEC using Dharmafect 1, and cells were either infected or RNA was collected 48 hrs post transfection.

V5-fused BPIFB3 was generated by cloning into pcDNA3.1/V5-His TOPO TA according to the manufacturer’s protocol. Development of GFP tagged RETREG1 and RETREG1mutLIR have been described elsewhere(22). RETERG1 GFP1-10 and BPIFB3 GFP11 were cloned into pcDNA3.1-GFP(1-10) or pEGFP-GFP11 respectively, using plasmids provided by Seema Lakdawala (University of Pittsburgh). RFP tagged LC3 cloning has been described previously(46). Plasmids were reverse transfected into U2OS cells using either X-tremeGENE 9 or X-tremeGENE HP according to the manufacturers protocol and fixed for fluorescence microscopy or infected at 48 hrs post transfection.

### RNA extraction, cDNA synthesis, and RT-qPCR

RNA was isolated using the GenElute Total RNA MiniPrep kit from Sigma according to the kit protocol. RNA was reverse transcribed using the iScript cDNA Synthesis kit (Bio-Rad) with 1 μg of RNA per sample. RT-qPCR was performed using IQ SYBR green SuperMix (Bio-Rad) in a Bio-Rad CFX96 Touch real-time PCR detection system. A modified threshold cycle (ΔCT) method was used to calculate gene expression using human actin for normalization. Primer sequences for actin, DENV, ZIKV, CVB, BPIFB3, and RETREG1 have been described previously(22,46).

### RNAseq

Total RNA was isolated as described above, and RNAseq was performed as previously described(25). Analysis of RNAseq data sets was performed using CLC Genomics 11 (Qiagen) to process and map sequences to the human genome (hg19) or the appropriate viral genome to calculate viral fragments per kilobase of transcript per million mapped reads (FPKM) values. Differentially expressed genes were identified using the DeSeq2 package in R with a significance cutoff of 0.001 and a fold change cutoff of two(47). Gene set enrichment analysis (GSEA) and manual sorting were used to identify pathways or specific transcripts differentially regulated. Generation of heat maps was done using MeViewer software based on ln(RPKM) values.

### Antibodies

Mouse monoclonal anti-V5 epitope tag was purchased from Invitrogen (R960-25). Rabbit polyclonal antibodies against CKAP4 (16686-1-AP), RTN4 (10950-1-AP), and FAM134B (21537-1-AP) were purchased from ProteinTech. Rabbit polyclonal antibodies to DENV NS3 (GTX124252) and ZIKV NS4B (GTX133311) were purchased from GeneTex. Recombinant mouse monoclonal anti-dsRNA was provided by Abraham Brass (University of Massachusetts). Alexa Fluor conjugated secondary antibodies were purchased from Invitrogen.

### Immunofluorescence and electron microscopy

Immunofluorescence microscopy was performed on cells grown in 8-well chamber slides (company?), fixed in 4% paraformaldehyde, and permeabilized with 0.1% Triton. In some cases, cells were fixed in ice cold methanol. Primary antibodies were incubated in PBS with cells for 1 hr, followed by staining with Alexa Fluor conjugated secondary antibodies for 30 min. Slides were mounted with coverslips using VectaShield containing 40-6-diamino-2-phenylindole (DAPI). Imaging was performed on an Olympus IX83 inverted microscope. All image quantification was performed using ImageJ/FIJI. Pixel intensity measurements were performed using isolated channels on individual cells with the region of interest (ROI) manager. Data are presented as mean pixel intensity, normalized to cell area. Quantification of fluorescent puncta was performed manually, counting the number ER localized vesicles alone, or co-localized with the indicated marker. Preparation of samples for TEM were done as previously described, by the Center for Biologic Imaging (University of Pittsburgh)(46). Imaging was performed on a JEOL 1011 transmission electron microscope. Quantification of TEM images was performed manually.

### Statistical analyses

All analyses were performed using GraphPad Prism. Experiments were performed at least three times. Student’s t test, 2way analysis of variance (ANOVA), or one-way ANOVA were used where indicated. Analysis of fluorescent microscopy data was done using a non-parametric Kruskal-Wallis test. Data are presented as mean ± standard deviation, with specific p-values detailed in the figure legends.

## Acknowledgements

We thank Abraham Brass (University of Massachusetts) for providing anti-dsRNA antibody, Theodore Pierson (NIH/NIAD) for providing the providing DENV replicon constructs, and Seema Lakdawala (University of Pittsburgh) for providing Split GFP constructs. This project was supported by NIH R01-AI081759 [C.B.C.]. In addition, C.B.C. is supported by a Burroughs Wellcome Investigators in the Pathogenesis of Infectious Disease Award.

